# Generation of the First Transgenic Line of the Iconic Coral Reef Fish *Amphiprion ocellaris*

**DOI:** 10.1101/2024.06.05.597662

**Authors:** Gabriel J. Graham, Emma M. Ibanez, Laurie J. Mitchell, Karen E. Weis, Lori T. Raetzman, Fabio Cortesi, Justin S. Rhodes

**Affiliations:** Department of Psychology, University of Illinois, Champaign, IL, USA; Department of Molecular & Cellular Biology, University of Illinois, Champaign, IL, USA; School of Biological Sciences, The University of Queensland, Brisbane, QLD, Australia; Department of Molecular & Integrative Physiology, University of Illinois, Champaign, IL, USA; Queensland Brain Institute, The University of Queensland, Brisbane, QLD, Australia; School of the Environment, The University of Queensland, Brisbane, QLD, Australia; Marine Eco-Evo-Devo Unit, Okinawa Institute of Science and Technology, Onna son, Okinawa 904-0495, Japan

**Keywords:** Anemonefish, Sex change, Transgenic, *Tol2*, Ef1a, Green Fluorescent Protein

## Abstract

The common clownfish, *Amphiprion ocellaris*, is an iconic coral reef fish, ubiquitous in the marine aquarium hobby and useful for studying a variety of biological processes (e.g., mutual symbiosis, ultraviolet vision, and protandrous sex change). Recently, CRISPR/Cas9 methods were developed for knocking out specific genes for mechanistic studies. Here, we expand the genetic toolkit for *A. ocellaris* by creating the first transgenic line using the *Tol2* transposon system. Fertilized eggs were co-injected with *Tol2* transposase mRNA and a plasmid encoding an Elongation factor 1 *α* (*Ef1α*): Green fluorescent protein (*GFP)* cassette at various concentrations, needle tip dimensions and timepoints post-fertilization. We compared various injection parameters and sterilization methods to maximize the survival of injected eggs. F0s (n=10) that were genotyped GFP+ were then raised to 6 months of age and crossed with wild-type (WT) females to confirm germline transmission. F1 offspring were also raised and crossed in the same manner. The highly efficient *Tol2* transposon system resulted in a 37% rate of transgenesis for surviving eggs amounting to a 2.7% yield of all injected eggs surviving and being GFP+ (n= 160). Of these, 10 were raised to adulthood, 8 spawned, and 5/8 (62.5 %) produced GFP+ offspring. Further, two F1s crossed with WT females produced 53.8% and 54.2% GFP+ offspring respectively, confirming the creation of a stable line. This is, to our knowledge, the first generation of a transgenic line in any coral reef fish. The ability to express transgenes of interest in the iconic anemonefish opens the door to a new era of exploration into their fascinating biology.

## Introduction

The false clown anemonefish *Amphiprion ocellaris* is a prominent icon of the coral reef. Anemonefish are famous for their mutualistic symbiosis with sea anemones, popularity in the marine aquarium hobby, and presence in popular culture. *Amphiprion* species also serve as important model organisms in the life sciences for studying a variety of biological processes such as larval recruitment, ultraviolet vision, monogamy, female behavioral dominance, protandrous sex change, paternal care, and mutualistic symbiosis with anemones (Beldade et al., 2016, Fricke and Fricke, 1977, Laudet and Ravasi, 2022, Mitchell et al., 2021, Buston and García, 2007, Mitchell et al., 2024). *A. ocellaris* live in small groups with a small home range immediately surrounding their host sea anemone. This ecological feature provides a number of logistical advantages for using anemonefish in laboratory and field research. In the laboratory, anemonefish are comfortably housed in modestly sized aquaria, and readily display natural physiology and behavior including protandrous sex change (Dodd et al., 2019, Parker et al., 2023, Parker et al., 2022, Madhu et al., 2010) and parental care (DeAngelis and Rhodes, 2016, Phillips et al., 2020, DeAngelis et al., 2017, DeAngelis et al., 2020, Ghosh et al., 2012, Barbasch et al., 2022). Additionally, larvae are easily reared in aquaria, meaning their entire life cycle can be studied under experimentally controlled conditions in the lab (Mitchell et al., 2021, Madhu et al., 2006, Roux et al., 2021, Laudet and Ravasi, 2022).

Recently, we developed methods for gene knockout in *A. ocellaris* using CRISPR/Cas9 technology (Mitchell et al., 2021). Briefly, we injected CRISPR/Cas9 reagents into *A. ocellaris* eggs, reared larvae bearing the null allele to adulthood, and genotyped them to confirm the gene had been knocked out. While these methods opened the door for genetic manipulation in a single anemonefish, we did not confirm that the mutation was present in the germ cells, and did not attempt to found mutant lines for longer-term studies.

Continued use of the anemonefish as a model organism will require methods for stable germ line transmission and methods for inserting genes into the genome. This includes inserting genes from other species or modified genes, also referred to as transgenesis. Transgenes can be selectively expressed in certain cells by using the promoter of a gene of interest to drive their expression. This approach is often used to visualize cell populations that express a gene by using the promoter of that gene to control the expression of a reporter such as green fluorescent protein (GFP) or lacZ (Ma et al., 2015). Other purposes include inserting genetic constructs that allow the conditional knockout or spatiotemporal control over expression of specific genes (e.g., cre-lox, doxycycline) (Campbell et al., 2012, Felker and Mosimann, 2016). Transgenes expressing channel rhodopsin (Seki et al., 2023), or designer chemoreceptors can be used to manipulate the activation of specific cell types (Silic and Zhang, 2021), and transgenes encoding designer calcium indicators (e.g., GCaMP) may be used to measure cellular activity (Muto et al., 2011). The diversity and capabilities of genetic tools available are expanding rapidly, thus it is important and timely to develop methods in anemonefish for transgene insertion.

While CRISPR/Cas9 technology has been used to insert genes by cut and replace, its reliability and efficiency can be low in certain cell types and organisms, making it challenging to achieve desired gene insertion without extensive screening and selection processes (Rozov et al., 2019). In contrast, the medaka (*Oryzias latipes*) derived transposon *Tol2* provides one of the most efficient methods to date for transgene insertion and is routinely used in zebrafish (*Danio rerio*) resulting in up to 50% of injected embryos containing the transgene in the germline (Kawakami, 2007). In addition, *Tol2* has been successfully adapted for use in several non-model organisms making it an ideal candidate for use in *A. ocellaris* (Fujimura and Kocher, 2011, Juntti et al., 2013, Stahl et al., 2019).

While methods for *Tol2*-mediated transgenesis have been established in other non-model organisms, attempting transgenic engineering in *A. ocellaris* brings new challenges, specifically regarding the microinjection of eggs laid on a solid substrate and high embryonic to larval mortality not found in other species (Yamanaka et al., 2021). Moreover, establishing stable transgenic lines that can be used for longer-term studies has never been accomplished in *A. ocellaris* or any coral reef fish to our knowledge. Here, we describe methods for the generation of the first stable transgenic line of anemonefish. As proof of principle, we used the *Tol2* transposon system to create several lines of *A. ocellaris* that express green fluorescent protein (GFP) under the control of the ubiquitous promoter for elongation factor-1 *α* (*Ef1α*). We describe methods for microinjection, artificial incubation, larval rearing, and transgene detection for future use in transgenic line development in *A. ocellaris*.

## Methods

### Animals and husbandry

Fish used in this study were either first or second-generation bred in-house from fish originally obtained from Ocean Reefs and Aquariums (Fort Pierce, FL), obtained as a gift from Peter Buston, or purchased from the pet trade. Fish were housed in 20-gallon tall (24” x 12” x 16”) or 25-gallon cube (18”×18” ×18”) centrally filtered aquariums. Salinity was kept at 1.026 specific gravity, temperature was kept between 26-28°C, pH was kept between 8.0 and 8.4, and lighting was provided by fluorescent and LED lights over tanks on a 12:12 light schedule (lights on at 0700 h and off at 1900). A 6” terra cotta pot served as a nest site and spawning substrate. Fish were fed twice daily with ReefNutrition TDO Chroma Boost pellets (Reed Mariculture Inc., Campbell, CA). Experimental procedures were approved by the University of Illinois Institutional Animal Care and Use Committee.

### Plasmids and *Tol2* RNA preparation

Plasmid DNA for pT2KXIGΔin (*Ef1α*:*GFP* expression vector) and pCS-ZT2TP (*Tol2* plasmid) was prepared using the Endo-Free Plasmid Maxi kit (Qiagen, Germantown, MD) according to the manufacturer’s instructions. To generate *Tol2* RNA, DNA from pCS-ZT2TP was digested overnight with NotI restriction enzyme using the FastDigest NotI system (Thermo Fisher Scientific, Waltham, MA). Linearized DNA was purified using the QiaQuick PCR purification kit (Qiagen) according to the manufacturer’s protocol. Approximately 3-5 µg of linearized plasmid DNA was used to generate capped *Tol2* RNA using the mMessage mMachine SP6 transcription kit (Invitrogen, Carlsbad, CA). In vitro transcribed RNA was purified using the RNeasy Mini kit (Qiagen) prior to use in injections.

### Preparation of eggs

In our lab, *A. ocellaris* pairs spawn every 13 to 23 days, typically in the afternoon hours between 16:00 and 19:00. On the day of spawning, anemonefish display characteristic behaviors and morphology of the genitalia. Females display a distended midsection and extended urogenital tube and the pair spends the majority of their time in the terra cotta pot. On the day of spawning, anemonefish clear the pot of algae, detritus, and microorganisms by vigorously biting and fanning with their pectoral and caudal fins. Immediately prior to spawning, females begin false egg laying runs, rubbing their ovipositor over the spawning site. Clutches typically consist of around 1000 eggs. Using these behavioral and morphological characteristics, we identified pairs that were going to spawn and waited until spawning was initiated as defined by the initiation of egg laying. *A. ocellaris* females attach the base of the egg to the surface of the terracotta pot, making a clutch of eggs look like a dense patch on a terracotta pot. The male will fertilize the eggs while the female is laying eggs. 15-30 minutes after the initiation of spawning, terra cotta pots containing eggs were removed from the aquarium and replaced with clean terra cotta pots. To prepare the eggs for microinjection, terra cotta pots were broken with a hammer and chisel until egg-containing tiles could fit into a petri dish. To keep track of injected eggs during the microinjection process, we removed eggs from the tile so that the remaining eggs were in neat rows with space between each row. Tiles were then kept in a small container of saltwater from the aquarium where the eggs were from prior to injection.

### Needle Preparation

A borosilicate glass micropipette (hereafter referred to as needle) (Harvard Apparatus: 1.0×0.58×100 mm; Harvard Apparatus, Holliston MA) was pulled using a Sutter Instrument P-97 (Sutter Instrument, Novato CA) using the following parameters: (Heat: 530, Pull: 150, Vel: 75, Time: 250). The tip of the needle was then broken using iris scissors under a light microscope, producing tip diameters between 5 and 15 µm. Needles with diameters outside this range were discarded.

### Transgene Insertion

Eggs were injected with a solution consisting of 1.4 µl of nuclease-free water, 0.7 µl of 2M KCl, 0.7 µl of phenol red, 1.2 µl of varied concentrations of *Tol2* mRNA, and 1.2 µl of the pT2KXIGΔin plasmid (Urasaki et al., 2006) containing the *Ef1α*:*GFP* cassette flanked by *Tol2* recognition sites at varied concentrations. The injection solution was backloaded into the needle using an Eppendorf microloader tip (Eppendorf catalog no. 930001007; Eppendorf AG, Hamburg, Germany), and the needle was loaded into a pneumatic microinjector (Narishige IM400; Narishige International, Amityville, NY). Air pressure and pulse duration were manipulated for each needle such that 0.75-1.25 nl of injection solution was dispensed per pulse, one pulse was delivered per egg.

Eggs were placed in a petri dish and immersed in Yamamoto’s ringer’s solution during injection to maintain osmotic homeostasis when the chorion is pierced by the needle during injection (Mitchell et al., 2021). Eggs were injected near the animal pole at the bottom of the yolk (see Fig. 1A in Mitchell et al., 2021). Injections took place at room temperature (∼23°C). After injection, eggs were counted and placed into artificial incubation.

**Figure. 1.**
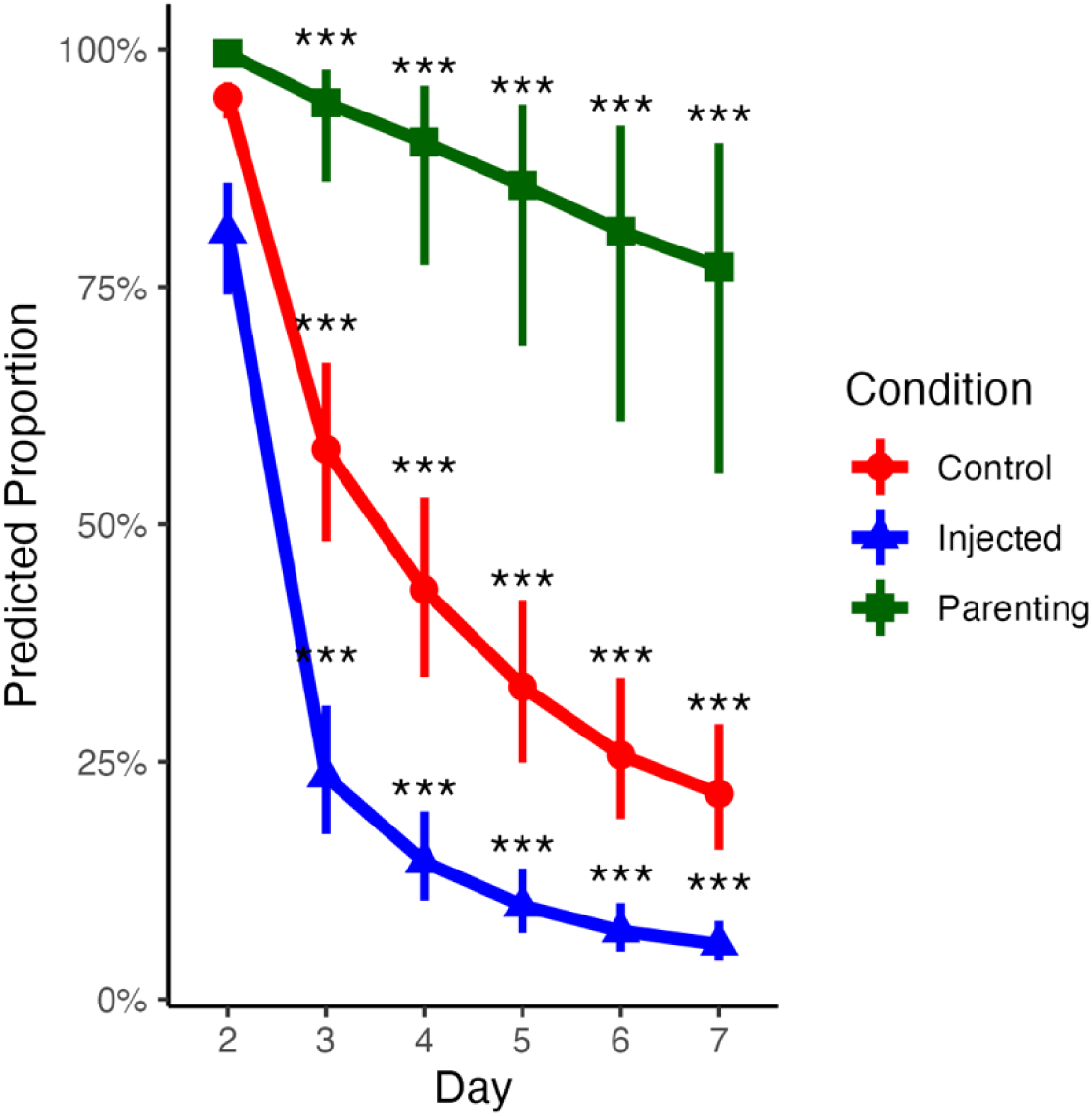
Glmer estimates for egg survival across dpf 1 – 6 for injected (blue triangle), control (red circle), and parented (green square) embryos. In all conditions, embryonic mortality increased significantly after 2 dpf. 0 Dpf was not included as the count on 0 dpf was used as the initial baseline for calculating survival. Dpf was fit as a categorical variable in these models to allow for non-linear relationships. Estimates are collapsed across all other covariates. “***” Indicates p<0.001, and vertical lines indicate 95% CI.

### Artificial Incubation

Tiles containing eggs were placed in 10-gallon tanks (20” x10” x12”) with plastic egg crating at the bottom acting as a platform for the tiles to lay above the floor of the tank. The tiles were arranged near wooden air stones with high amounts of airflow to maintain a moderate level of agitation and aeration.

### Experiment 1: Injection Parameters

To identify variables that impact egg survival and transgenesis rate, we tracked needle tip diameter, *Ef1α:GFP* plasmid concentration, *Tol2* mRNA concentration, and time between initiation of spawning and removal of eggs (time to inject).

### Experiment 2: Sterilization Methods

#### Tank Sterilization

We compared two tank sterilization methods to reduce embryonic mortality caused by disease or infection: adding 0.7 ml of 2.3% methylene blue (Kordon catalog no. 37344, Kordon Methylene Blue Disease Preventative; Kordon, Hayward, CA) to the tank water, or continuous ultraviolet (UV) sterilization of the water using a UV sterilizer (Aqua Ultraviolet catalog no. 617750002859; Aqua Ultraviolet, Temecula, CA). It was not possible to use both methods simultaneously because UV sterilization breaks down methylene blue (Huang et al., 2013). Tiles that had injected eggs as well as control tiles with uninjected eggs were included. Dead embryos were removed once a day until hatching, while surviving embryos were quantified until 6 days post fertilization (dpf) where day 0 is the day of fertilization. To compare the efficacy of artificial incubation to parental incubation, surviving embryos under parental care were also quantified daily.

#### Dips

In addition to UV sterilization, tiles were treated daily with different chemicals dissolved in freshly made 35 ppt saltwater. Specifically we tested: 1) 0.01% Povidone iodine in saltwater for 10 minutes (Park et al., 2019) , 2) 0.005% methylene blue in saltwater for 10 seconds (Park et al., 2019), 3) 0.15% KCl in saltwater for 5 minutes (Davis et al., 2018), or 4) 0.12% H_2_O_2_ in saltwater for 5 minutes (Wagner et al., 2012). In addition, we tested a freshwater dip for 3 minutes (Robinson et al., 2008). Concentrations and contact times of the dips were reduced from the source protocols due to embryos’ increased sensitivity to chemical exposure relative to adults (Mohammed, 2013). Dead embryos were removed from the tiles before dips and tiles were rinsed in clean saltwater after each dip before being returned to incubation tanks. Dips were performed daily until 6 dpf.

### GFP Detection

#### Fluorescence Microscopy

Embryos 6 dpf were viewed under 485 nm fluorescence microscopy (Zeiss SteREO Discovery V20) to screen for GFP expression. Any embryos that didn’t display fluorescence were removed prior to hatching.

### Hatching

On day 7 post fertilization, tiles were mounted vertically with mild aeration in 10-gallon hatching tanks with all sides blacked out to prevent intrusion of lateral light. While some transgenic larvae were able to hatch with this method, we encountered a similar problem as was described in Mitchell et al., (2021) where injected embryos would struggle to hatch and eventually die inside the chorion by 11 dpf. Therefore, embryos that failed to hatch by 8 dpf were removed from the tiles and were manually hatched as described in Mitchell et al. (2021). We used fine-tipped forceps to grasp the bottom of the chorion near the larval tail and dissection scissors were used to make an incision in the bottom of the chorion. Then we used a second pair of fine-tipped forceps to grasp the cut in the chorion and tear it open, freeing the larva (Mitchell et al., 2021). All larvae were then returned to the hatching tanks along with a low flow sponge filter. Larvae were fed live rotifers (*Brachionus* spp.) at a concentration of ∼10 rotifers/ml. 300 µL of cryopreserved *Nanochloropsis* spp. (Nanochloropsis Cryo-Preserved Algae Paste; Brine Shrimp Direct, Ogden, UT) was added to the water daily to feed the rotifers.

### Rearing

Larvae were fed live rotifers until 4 days post hatch (dph) where 0 dph is the day of hatching. At 5 dph, larvae were transitioned to live *Artemia* nauplii and TDO Chroma Boost A pellets (Reed Mariculture, Campbell, CA). Around 13 dph, larvae began metamorphosis. During their larval stage, *A. ocellaris* spent most of their time at the surface of aquariums near walls eating rotifers and *Artemia*, however, once metamorphosis began, juvenile fish retreated to dark areas in the bottom of aquaria and decreased overall movement. Metamorphosis lasted around 14 days after which juveniles were fed TDO Chroma Boost B pellets (Reed Mariculture, Campbell, CA) and continued to be fed live *Artemia* nauplii.

#### PCR

A small fin clip was taken from the tail fin of juveniles when they were approximately 2.5-3.0 cm long (4-6 months old). Genomic DNA was extracted using a Qiagen DNeasy Blood & Tissue Kit (Qiagen catalog no. 69504; Qiagen, Valencia, CA). PCR amplification was performed using primers targeting the GFP coding sequence, primers targeting *GnRH1* were used as a positive control, primers with no DNA were used as a negative control. One*Taq*^®^ Hot Start Quick-Load^®^ 2X Master Mix with Standard Buffer (New England Biolabs catalog no. M0488S; New England BioLabs, Ipswich, MA), was used in a 25 µl reaction with 2 µl template DNA for amplification according to the manufacturer’s guidelines. We performed gel electrophoresis on a 2% agarose gel with 10 µl 10 mg/mL ethidium bromide (EtBr) submerged in 1.5L 1X TBE with 150 µl 10 mg/mL EtBr (Sigma-Aldrich cat no. E1510; Merck KGaA, Darmstadt, Germany). Wells in the gel were loaded with 10 µl amplified DNA, along with a well containing 10 µl Quick-Load^®^ Purple 1 kb DNA Ladder (New England Biolabs catalog no. N0552S; New England BioLabs, Ipswich, MA) and ran at 105 V for 45 minutes. Bands were analyzed using a Gel-Doc EZ Imager (Bio-Rad, Hercules, Ca) set to low band detection sensitivity.

### Germline Transmission

GFP+ individuals identified via PCR genotyping (F0) were raised to 6 months of age before being paired with a wild-type (WT) female. Eggs fertilized by these F0 founders were then observed under 485 nm fluorescence microscopy at 6 dpf and the number of their embryos (F1) expressing GFP were quantified (see table 1 for success rate). GFP+ F1 embryos were hatched and subsequently raised to 6 months before being genotyped as described above. Confirmed GFP+ F1s were then crossed with WT females to evaluate the proportion of GFP+ F2 embryos.

**Table 1.** Rate of germline transmission produced by GFP+ F0s crossed with WT females.

| F0 x WT | % Embryos GFP+ |
| --- | --- |
| TAM | 6.725 |
| R1T | 4.918 |
| R2B | 4.711 |
| R8TB | 1.613 |
| R3T | 1.222 |
| B15M | 0 |
| R4T | 0 |
| R5T | 0 |

### Statistical Analyses

R (4.2.3) was used for statistical analyses with a p<0.05 considered statistically significant. The glmer() function from the R package ‘lme4’ was used to fit mixed logistic regression models predicting the proportion of embryos surviving on a particular day as a function of various factors and covariates. The response variable (i.e. proportion of embryos surviving) was measured on dpf 0-6 with the count on dpf 0 defining 100%. The “family= binomial(logit)” option was used within the glmer() function to model the binomial error structure. Because tiles were taken from batches of eggs, and batches of eggs were taken from individual spawning pairs, we used models that included a nested random effect where tile was nested within batch which was nested within parents to account for the repeated measures structure and inevitable correlation between days as eggs disappear from the tiles. Dpf was always entered as a factor instead of a continuous variable to allow for non-linear relations. Other fixed effects depended on the experiment, and included factors such as tank sterilization method, dip, and continuous covariates such as needle tip diameter, *Tol2* mRNA concentration, *Ef1α*:*GFP* plasmid concentration, time to inject eggs, and time to take eggs. Injected eggs and control eggs were evaluated in separate models due to the various terms only present in the injected condition such as the needle tip diameter. We calculated 95% confidence intervals (CI), odds ratios (OR), and Post-hoc tests using the package “emmeans”, and anovas were calculated using the Anova() function from the package “car” with type III tests specified.

## Results

In total, we injected 6,017 eggs on 107 tiles created from 45 batches laid by 20 different spawning pairs. We also artificially incubated 5,013 uninjected eggs until the point of hatching from each of the injected batches using the same artificial incubation conditions to serve as controls. For comparison, we also tracked the survival of 5,049 eggs naturally reared by parents to establish a best-case scenario for embryonic survival. As days progressed, embryos were lost due to lack of fertilization, infection, or removal by the parents (in naturally reared controls). This was indicated by a significant effect of dpf in the mixed logistic regression (p<0.001; see Supplementary table 1A for ORs and 95% CIs for each dpf). However, the loss was greater for injected or artificially reared control embryos (Fig. 1) as indicated by a significant effect of condition in the mixed logistic regression (n = 16079, χ^2^ = 211.09, p<0.001). Naturally reared embryos had significantly better survival than artificially incubated embryos (p<0.001, OR = 0.08, 95% CI [0.03, 0.23]) and injected embryos (p<0.001, OR = 0.02, 95% CI [0.01, 0.01]), and injected embryos had significantly worse survival than artificially reared controls (p<0.001, OR = 4.49, 95% CI [3.57, 5.63]) (see Supplementary table 1A for the full model summary and Supplementary table 1B for pairwise comparisons). Overall, we found that approximately 8.8% (n = 424/4820, SE = ± 0.002) of injected embryos survived until hatching as compared to 39.3% of control embryos (n = 1576/4013, SE = ± 0.006). Of the surviving injected embryos, 36.8% (n = 138/375, SE = ± 0.062) were identified GFP+ by fluorescence microscopy, and 55.6% (n = 10/18, SE = ± 11.712) of these GFP+ larvae genotyped GFP+ in a tail fin clip as juveniles. Of these 10 GFP+ F0s, 8 spawned, and 62.5% (n =5/8, SE = *±* 17.116) produced a fraction of F1 offspring carrying the transgene (see table 1 for GFP+ f1s produced by F0s). 53.8% (n = 14/26, SE = ± 8.268) of F2 larvae were GFP+.

### Experiment 1: Injection Parameters

#### Needle Diameter

As discussed in Mitchell et al., (2021), the diameter of the needle tip plays a significant role in egg survival after injection. When the needle tip diameter is too small (<5 µm), the needle has a tendency to bend and may even break on contact with the chorion. Additionally, smaller needle tips tend to clog easily or dispense less liquid than is required for transgenesis. Conversely, when the needle tip diameter is too large (>15 µm), the injection causes significant damage to the egg and results in high mortality. We therefore sought to find the optimal needle diameter to maximize the survival of transgenic embryos. Analysis of embryonic survival with needle diameter as a continuous variable indicated a significant negative effect on survival rate (n = 6017, χ^2^ = 10.996, p <0.001, OR = 0.86, 95% CI [0.79, 0.94]) (see Supplementary table 2 for the full model summary) (Fig. 2A). When the needle tip was between 5 and 9 µm, survival 6 dpf was approximately 11% (n = 331/2997, SE = ± 0.003) whereas when the needle tip was between 10 and 15 µm, survival dropped to 4.5% (n = 49/1085, SE = ± 0.004). In all subsequent analyses of survival rates of injected embryos below, needle tip diameter was included as a variable to account for this source of variation (see Supplementary tables 2,3, and 4).

**Figure. 2.**
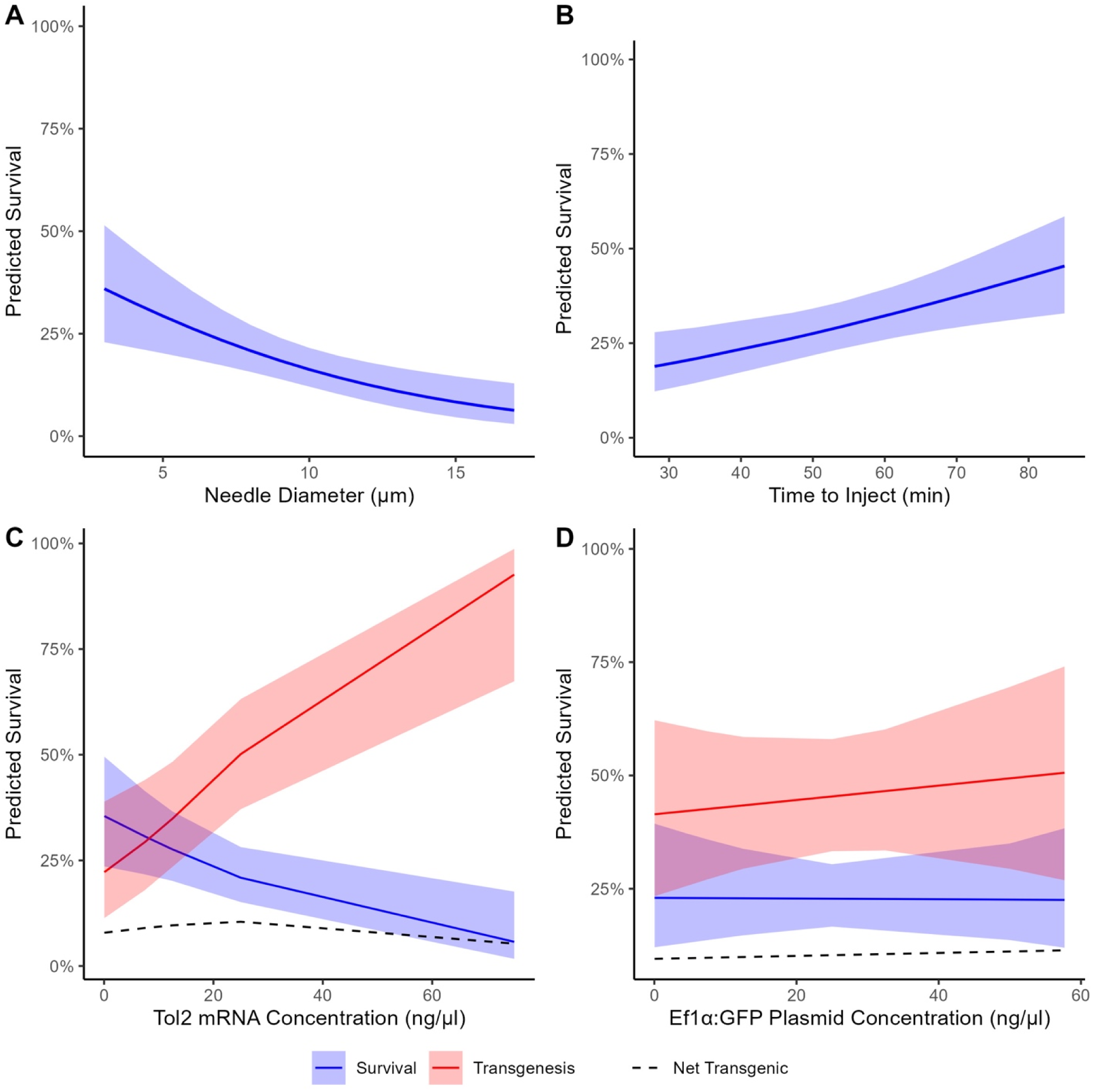
Glmer estimates for A) The survival rate of injected embryos by needle diameter. B) The survival rate of injected embryos by time to inject. C) The survival rate (blue), transgenesis rate (red), and the net yield of transgenic embryos (the predicted proportion of injected eggs surviving and being transgenic [black]) by the concentration of *Tol2* mRNA injected. D) The survival rate (blue), transgenesis rate (red), and the net yield of transgenic eggs (black) by the concentration of *Ef1α*:*GFP* plasmid injected. In all of these plots, estimates are collapsed across all other covariates. Line indicates estimates and ribbon indicates 95% CI.

#### Reagent Concentrations

When using a transposon to randomly insert a gene into the genome it is preferable to use the lowest possible plasmid and transposon concentrations to minimize mortality from cytotoxicity while still maintaining successful transgenesis. Thus, we manipulated the concentrations of *Tol2* mRNA and *Ef1α*:*GFP* plasmid in each injection with the goal of finding the concentrations that result in the highest yield of transgenic embryos. Previous studies using the *Tol2* transposon system co-injected 25 ng/µl of the transposase mRNA and 25 ng/µl of the plasmid DNA to successfully create transgenic lines, however, due to the high mortality we observed in *A. ocellaris,* we considered whether other combinations of reagent concentrations may be better for our organism (Stahl et al., 2019, Juntti et al., 2013, Detrich III et al., 2009, Fujimura and Kocher, 2011, Thummel et al., 2006). We therefore compared various combinations of *Ef1α*:*GFP* plasmid (0.058 ng/µl - 57.692 ng/µl) and *Tol2* mRNA (0.058 ng/µl – 75 ng/µl) concentrations injected into eggs to determine the impact on embryonic survival. *Ef1α*:*GFP* plasmid concentration had no significant effect on the survival of injected embryos (n = 2960, p = 0.102, χ^2^ = 2.680) (Fig. 2D) while *Tol2* mRNA concentration had a significantly negative effect on survival (n = 2960, p = 0.015, χ^2^ = 5.903, OR = 0.94, 95% CI [0.90, 0.99]) (Fig. 2C). We also found that there was no significant effect of the interaction between Ef*1α*:*GFP* plasmid and *Tol2* mRNA concentration (n = 2960, χ^2^ = 3.682, p = 0.055) (see Supplementary table 3 for the full model summary).

#### Time After Initiation of Spawning to Inject Eggs

Eggs were removed from the parents 15-30 minutes after the initiation of spawning to allow ample time for fertilization and for a sizeable yield to be obtained. Eggs were injected from 30 to 80 minutes after initiation of spawning. At 60-80 minutes post-fertilization, the chorion of the egg thickens, which prevents further injection of the egg with the needle (Mitchell et al., 2021). We recorded the amount of time between onset of spawning and egg injections and found a significant positive effect on survival (n = 1041, χ^2^ = 7.093, p = 0.008, OR = 1.02, 95% CI [1.01, 1.03]) (Fig. 2B) (see Supplementary table 4 for the full model summary).

We considered the possibility that breaking the terra cotta pots with a hammer and chisel during a period of time when the eggs are sensitive to damage could cause the positive correlation between survival and time until injection. However, this hypothesis was not supported because we found that there was no effect of time after the initiation of spawning to take eggs on survival of control eggs (n = 1871, χ^2^ = 2.544, p = 0.111) (see Supplementary table 5).

### Experiment 2: Sterilization Methods

#### Tank Sterilization

Preliminary experiments found that injected eggs could not be returned to parents because the fish will remove and eat injected eggs, suggesting that the parents can detect damage to the injection site from the needle. Parental care is important for egg survival because the father will agitate the eggs to promote oxygenation and clean the outer surface of individual eggs. Thus, we attempted to replicate behaviors involved in parental care within an artificial incubation set-up, namely, to keep the eggs oxygenated and protect them from infection during development.

Previously, we employed a common artificial egg incubation technique where air stones vigorously disturbed water in the tank and methylene blue acted as a decontamination agent in the water (Mitchell et al., 2021). In our case, this system was suboptimal because artificially incubated eggs had significantly lower survival rate than those incubated by their parents (Fig. 1, see above). Thus, we hypothesized that UV sterilization would perform better than sterilization via methylene blue. For both injected and control eggs, there was no significant difference between UV and methylene blue treatment (p = 0.305, χ^2^_1_ = 1.052), (χ^2^_1_ = 1.049, p = 0.306) (see Supplementary table 6 for the full model summary for control and Supplementary table 2 for full model summary the full model summary for injected eggs) respectively (Fig. 3A). Due to convenience, we used UV sterilization in all subsequent experiments.

**Figure. 3.**
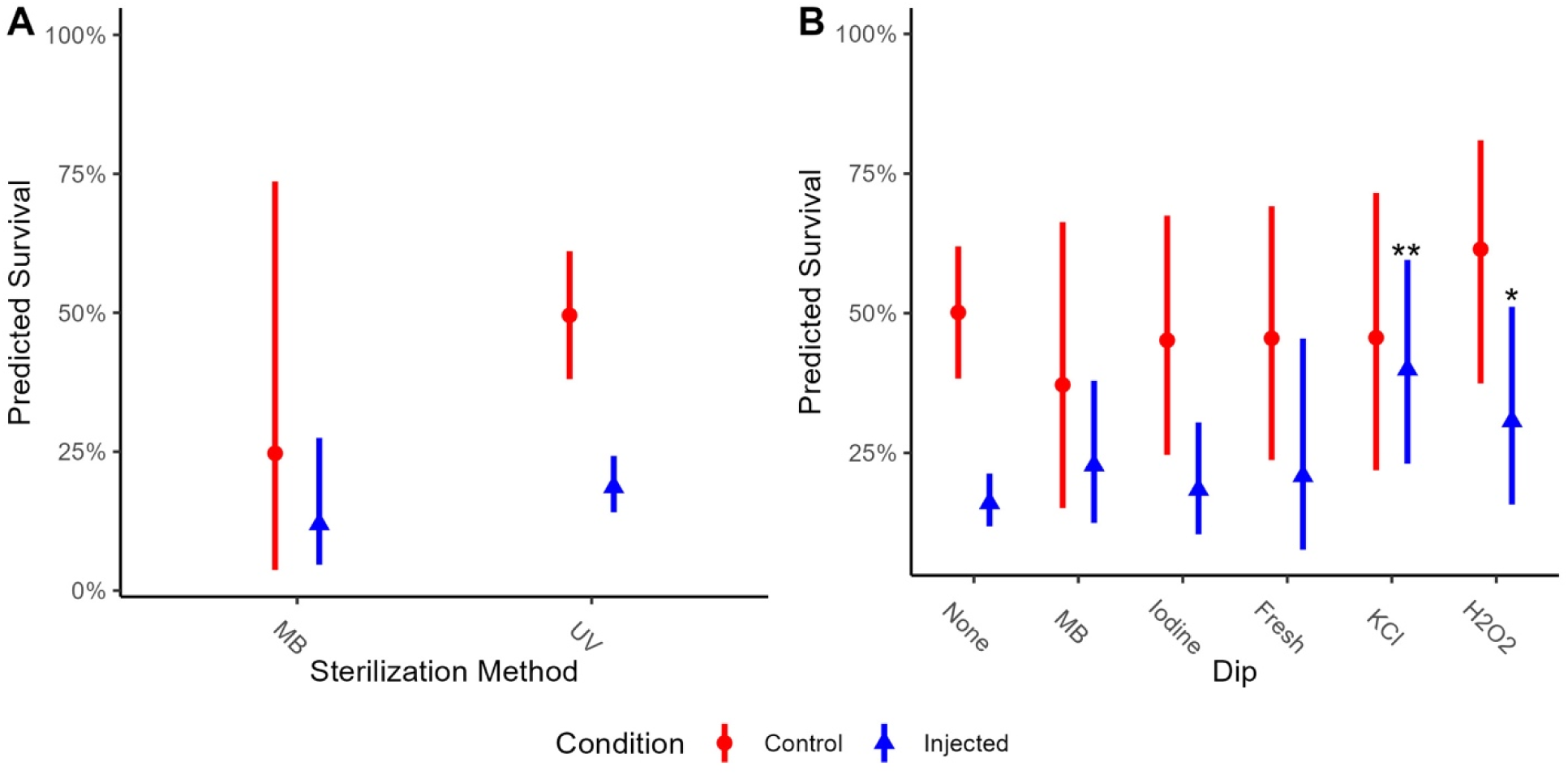
Glmer estimates for A) The survival rate of injected and control embryos by methylene blue (MB) or ultraviolet (UV) sterilization in control (red circle) and injected (blue triangle) embryos. There was no significant difference in survival when using MB sterilization compared to UV sterilization. B) The survival rate of injected (blue triangle) and control (red circle) embryos by no dip (None), 0.005% methylene blue in saltwater for 10 seconds (MB), 0.01% povidone iodine in saltwater for 10 minutes (Iodine), freshwater for 3 minutes (Fresh), 0.15% KCl in saltwater for 5 minutes (KCl) or 0.12% H_2_O_2_ in saltwater for 5 minutes (H_2_O_2_). For injected embryos, H_2_O_2_ and KCl are significantly better than no dip. Estimates are collapsed across all other covariates. Points indicate estimate, lines indicate 95% CI. “**” Indicates p<0.01.

#### Dips

We noticed that under both of the sterilization methods described above, there were still small arthropods and microorganisms living on the tiles and embryos suggesting the need for improved sterilization. In order to determine whether we could improve survival of embryos under artificial incubation, we compared their survival after dipping the tiles in different disinfectant solutions common in aquaculture or the aquarium hobby for short periods of time daily. For injected embryos, we found that two of the dips (H_2_O_2_ and KCl) resulted in significantly better survival than no dip (n = 6017, χ^2^ = 12.876, p = 0.025): H O (n = 90, p = 0.044, OR = 2.32, 95% CI [1.02, 5.28]) and KCl (n = 186, p = 0.001, OR = 3.49, 95% CI [1.62, 7.50]) (see Supplementary table 2 for the full model summary) (Fig. 3B). For control embryos, we found no significant effect of dip (n = 4615, χ^2^ = 3.349, p = 0.646) (see Supplementary table 6 for the full model summary) (Fig. 3B).

### Rate of Transgenesis

We then examined the effects of the concentration of *Ef1α*:*GFP* plasmid, *Tol2* mRNA, and their interaction on the rate of transgenesis for injected embryos surviving to 6 dpf. GFP+ embryos were quantified using fluorescence microscopy and compared against the total number of embryos remaining on that tile. *Ef1α*:*GFP* plasmid concentration did not have a significant effect on the rate of transgenesis (n = 375, χ^2^ = 3.039, p = 0.081) (Fig. 2D), while the *Tol2* mRNA concentration had a significantly positive effect (n = 375, p = 0.010, χ^2^ = 6.610, OR = 1.08, 95% CI [1.02, 1.14]) (Fig. 2C). The interaction between *Ef1α*:*GFP* plasmid and *Tol2* mRNA concentrations was not significant (n = 375, χ^2^ = 1.592, p = 0.207) (see Supplementary table 7 for full model summary).

### GFP expression pattern in F0s

GFP expression patterns in F0s were highly variable, with many expressing GFP only around the yolk, some expressing GFP only in the somatic tissue, and some expressing GFP in both somatic tissue and around the yolk (Fig.4). The various GFP expression patterns were not quantified.

**Figure. 4.**
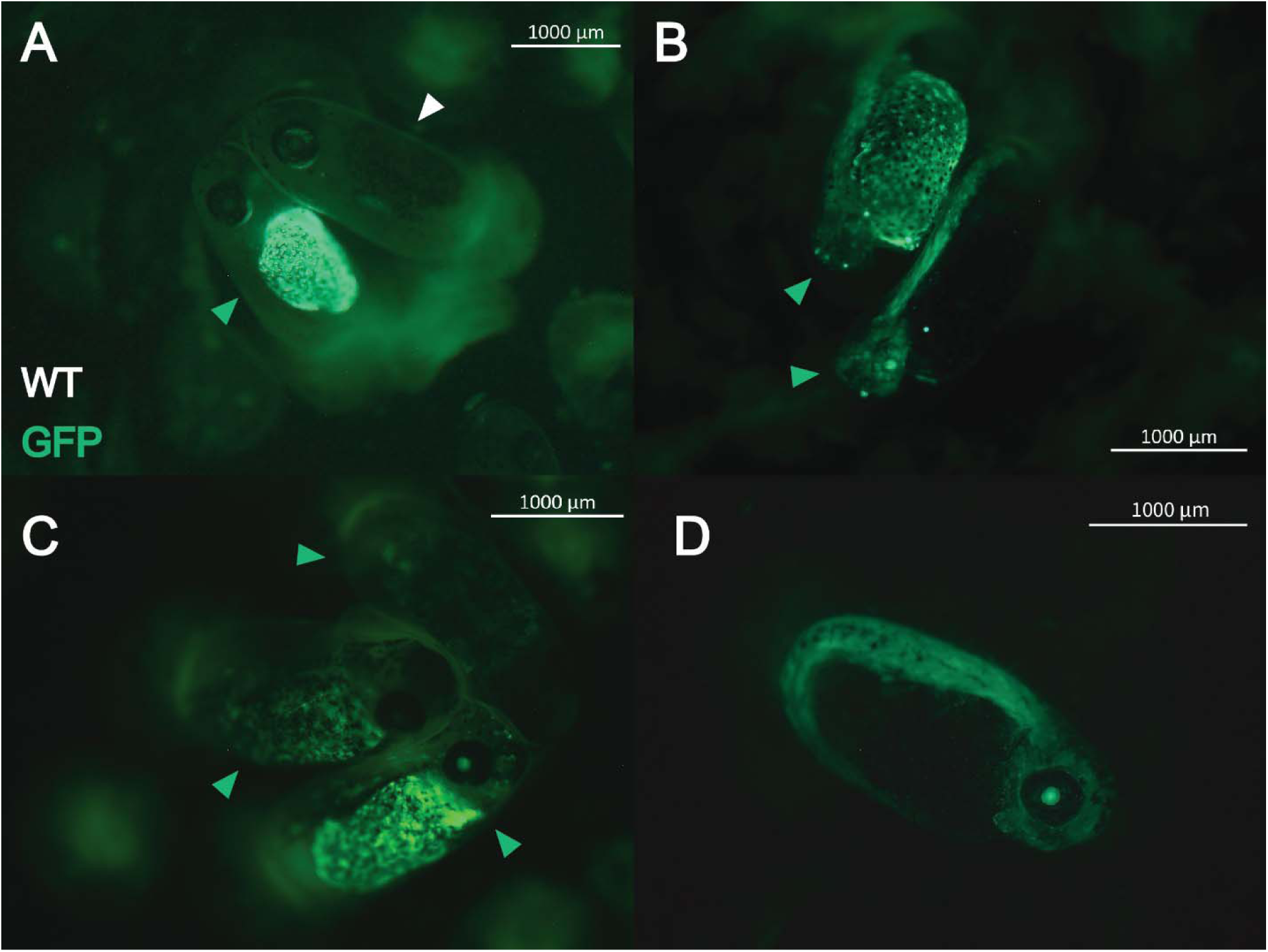
GFP+ and WT F0 *Amphiprion ocellaris* embryos. A) A WT embryo next to a GFP+ embryo with expression restricted to only the tissues around the yolk. Both embryos are at 6 dpf. B) Two F0s at 3 dpf showing differing GFP expression patterns, with the top expressing GFP both around the yolk and in the embryo proper and the bottom only expressing GFP in the embryo proper. C) Three GFP+ embryos at 6 dpf showing varying levels of mosaicism. The bottom shows the highest level of GFP expression, the second from the bottom showing a moderate level of GFP expression, and the top showing the lowest level of GFP expression. These embryos express GFP in both embryonic and extraembryonic tissues. D) An embryo at 6 dpf expressing GFP only in its embryonic tissues.

### Crossing and Progeny

Fluorescent embryos were hatched, raised to adulthood, and genotyped from a tail fin sample. Of these, 55.6% (n = 10/18, SE = ± 11.712) were GFP+ (Supplementary Fig. 1). These 10 GFP+ F0s were then paired with WT females at 6 months post-hatch. Of these, 8 spawned, and 62.5% (n = 5/8, SE = ± 17.116) produced GFP+ progeny as determined by fluorescence microscopy (Fig. 5) indicating successful transmission through the germline. The estimated fraction of F1 offspring that were transgenic for each F0 is shown in table 1. The mean proportion of GFP+ embryos produced by the 5 F0s that produced GFP+ offspring was 3.838% (SE = ± 1.053). We then raised 6 of these GFP+ F1s to adulthood, verified they were GFP+ via a tail fin sample (Supplementary Fig. 2. Two of these F1s spawned and produced 53.8% (n =14/28, SE = ± 8.268) and 54.2% (n = 26/48, SE = ± 7.192) GFP+ F2 progeny respectively, confirming line stability.

**Figure. 5.**
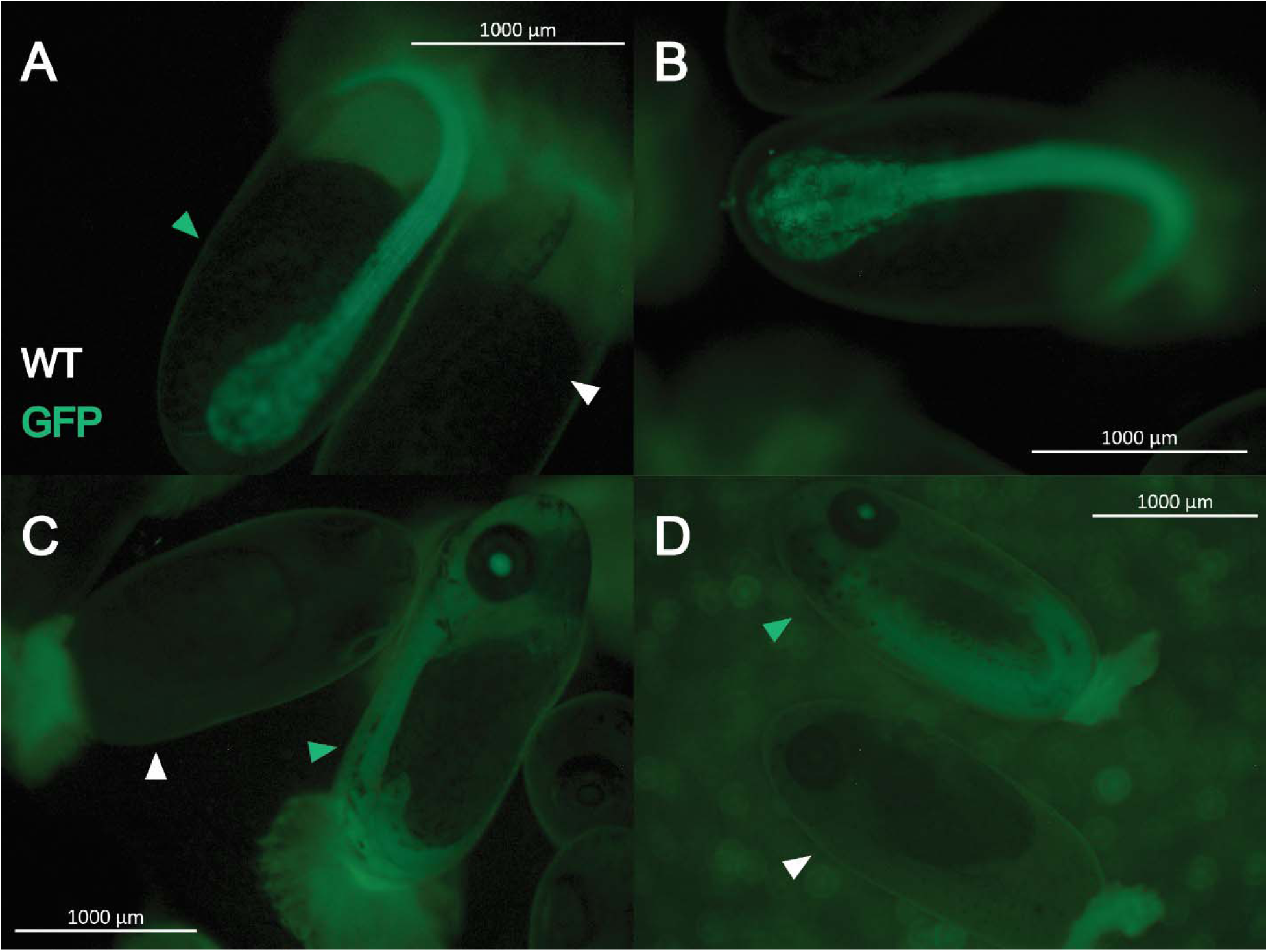
GFP+ and WT F1 *Amphiprion ocellaris* embryos. A and B) Show GFP+ embryos at 2 dpf. C and D) Show WT and GFP+ embryos at 6 dpf. All F1s display ubiquitous GFP expression in embryonic tissues and no yolk expression was detected.

## Discussion

We used the *Tol2* transposon system to generate the first transgenic line of *A. ocellaris* and, to our knowledge, the first transgenic line of any coral reef fish. While only 2.7% (n = 160/6017, SE = ± 0.206) of all injected embryos were both GFP+ and survived until hatching, the large size of *A. ocellaris* clutches (∼1000 eggs) means transgenic lines can be generated with relatively few spawns. Embryos produced from wildtype females crossed with GFP+ (F0) males produced on average 3.838% GFP+ F1 offspring. We confirmed these GFP+ F1 offspring are ubiquitously hemizygous for the transgene since approximately 50% of their progeny with a wildtype female were GFP+. Importantly, surviving F0s and F1s did not display any major phenotypic abnormalities, and developed as normal anemonefish. Taken together, this shows that *Tol2*-mediated transgenesis is an effective tool for creating stable transgenic lines of anemonefish.

A main goal of this study was to establish efficient methods for producing transgenic lines of anemonefish using the *Tol2* transposon system. Thus, we recorded factors that varied between injections including the identity of the breeding pair that laid the eggs, the diameter of the needle tip, and time after spawning when eggs were injected. Further, we manipulated the concentrations of *Ef1α:GFP* plasmid and *Tol2* mRNA and evaluated new sterilization procedures for artificial incubation. We found that implementing the following parameters resulted in the highest possible survival rate for injected eggs while maintaining efficacy for transgenic line production:

1. Using a needle with a tip diameter between 5 and 9 µm. Needle tip diameters smaller than 5 µm struggle to deliver the desired amount of liquid to the egg and struggle to pierce the chorion while needle tip diameters larger than 10 µm lead to excessive mortality.
2. Begin injecting eggs 40-50 minutes after the initiation of spawning. While survival increases with time to begin injecting, we recommend waiting 40-50 minutes after the initiation of spawning to begin injecting eggs in order to maximize the number of eggs that can be injected prior to chorion hardening. Longer wait times will drastically decrease yield while injecting too early results in significant death and should be avoided. This time window corresponds to taking eggs from parents around 30 minutes after the initiation of spawning due to the time taken to break the terra cotta pot into tiles and make rows of eggs. We injected eggs at a temperature of ∼23°C; increases in temperature will shift this window closer to the time of spawning while decreases will shift this window farther from the time of spawning.
3. UV sterilization during artificial rearing. We found no significant difference between UV sterilization and methylene blue sterilization in terms of embryonic survival, therefore we suggest using UV sterilization as opposed to methylene blue for reasons of convenience. Unlike UV sterilization, methylene blue requires repeat dosing over time to maintain effectiveness. However, methylene blue is cheaper and easier to acquire, so for few injections, methylene blue may be preferable.
4. Using a KCl dip. The KCl dip significantly improved survival rate compared to no dip for injected embryos while having no effect on control embryonic survival. This may suggest that KCl aids in sanitizing the wound left from injection or prevents infection caused by chorion damage.
5. Of the concentrations we tested, we suggest using a transgene plasmid concentration of 57.692 ng/µl and a *Tol2* mRNA concentration of 25 ng/µl. We tested the significance of *Tol2* mRNA concentration and *Ef1α*:*GFP* plasmid concentration and their interaction on both survival and rate of transgenesis. We found that in both cases, only the *Tol2* mRNA concentration had a significant effect. Based on the predicted proportion of embryos that survive and are transgenic (‘Net Transgenic’ in Fig. 2C), a concentration of 25 ng/µl produced the highest net transgenic rate (10.48%). Since the *Ef1α*:*GFP* plasmid concentration did not have a significant effect on survival or transgenesis, we suggest using a high concentration relative to the *Tol2* concentration to ensure that the transgene is present in sufficient concentration to interact with the *Tol2* transposase and be inserted into the genome. The highest concentration we tested was 57.692 ng/µl, though, in theory, higher concentrations would likely result in a high rate of transgenesis without having a significant impact on survival.

While a mosaic expression pattern was expected since transgenesis was anticipated to occur after the one-cell stage, the yolk-only versus embryo-only expression was surprising (Fig. 4). One possible explanation for yolk-only expression is accidental injection of transgene reagents into a part of the yolk more distal to the animal pole than optimal resulting in a relatively higher rate of transgenesis in the yolk syncytial layer (YSL) following the fusion of the cytoplasm of the cells that make up the YSL with the yolk (Gilbert, 2000, Carvalho and Heisenberg, 2010). While no eggs were injected after the one-cell stage, the time taken for injected *Tol2* mRNA to be properly translated and fold into a functional protein likely means that in mosaic embryos, the *Ef1α*:*GFP* cassette was inserted into the genome after one or several divisions. This problem may be reduced by injecting *Tol2* protein as opposed to *Tol2* mRNA as the protein does not have this translational delay and in many cases can integrate the transgene prior to the first division, though we did not try this in the current study (Ni et al., 2016).

The major limiting factor we experienced in the generation of transgenic lines was the low survival of embryos raised in artificial incubation. Control embryos displayed significantly lower survival than eggs reared by parents, and injected embryos displayed significantly lower survival than controls. We tried improving embryonic mortality in artificial incubation by using sterilizing dips and saw a moderate improvement in survival of injected embryos when using a KCl dip. Still, we feel more experiments varying KCl dip time and concentration are required to further improve sterilization of injected embryos. Mortality may also be reduced via substrate detached incubation as the number of pests introduced by tiles will be reduced, however this method has significant drawbacks as it requires removing eggs from their substrate which may damage them (Yamanaka et al., 2021). We also manipulated injection parameters such as needle tip diameter, *Ef1α:GFP* plasmid concentration, *Tol2* mRNA concentration, and time to begin injection in order to reduce the mortality of injected eggs. While several of these factors had significant effects on embryonic mortality, they are limited in the extent to which they can be varied while maintaining transgenesis.

The creation of methods for establishing transgenic lines of *A. ocellaris* has significantly increased the power of *A. ocellaris* and other closely related anemonefish as laboratory model species. The majority of research done on anemonefish takes place in the wild despite anemonefish being remarkably well suited to laboratory conditions (Laudet and Ravasi, 2022). In the near future, we are interested in generating transgenic anemonefish that allow manipulation of specific neuronal and radial glial cell populations either via insertion of chemogenetic or optogenetic receptors, or by inserting endogenous genes under alternative promoters to examine how they impact the rate or manner of sex change. Additionally, neuronal and glial cell populations that regulate sex change could be visualized by expressing GFP in these cells. Anemonefish are increasingly being used to study multiple interesting biological phenomena related to monogamy, female behavioral dominance, territorial aggression, paternal care, larval recruitment, color vision, and symbiosis with sea anemones (Laudet and Ravasi, 2022). In the context of exploring these biological questions and others, the experimental power associated with the capability to generate transgenic lines of anemonefish is nearly endless.

## Supplementary Information

Supplementary material is available.

## Author Contribution

Experimental design: JSR, LJM, GJG. Performed the experiment: GJG, EMI. Analyzed the data: GJG, JSR. Provided Materials: JSR, LTR, KEW. Wrote the manuscript: GJG, JSR.

## Data availability

Data will be made available on request.

## Declarations

### Ethics Approval

*A. ocellaris* is not a protected or endangered species. All experimental procedures were approved by the University of Illinois Institutional Animal Care and Use Committee.

