## supplementary Fig. for "Generation of the First Transgenic Line of the Iconic Coral Reef Fish *Amphiprion ocellaris*"

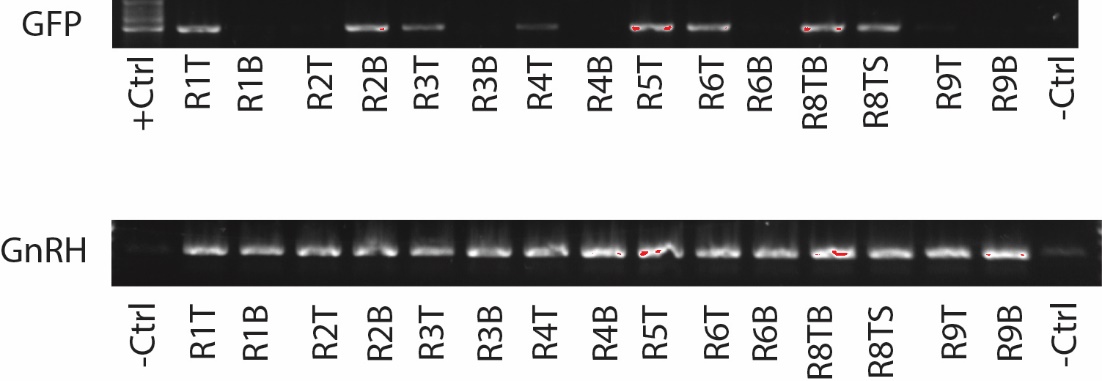

**Supplementary Figure.1** Genotyping of 8/15 GFP+ individuals (top) and GnRH control (bottom)

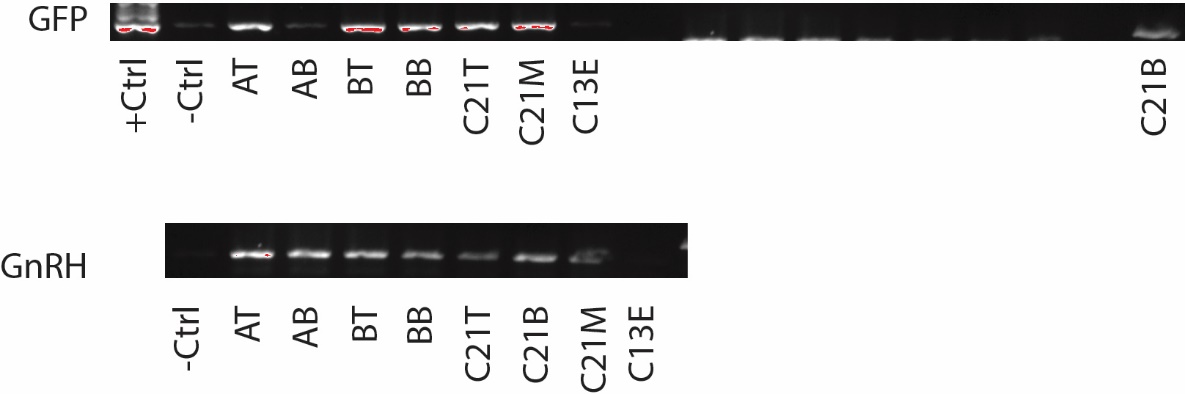

**Supplementary Figure. 2** Genotyping of GFP+ F1 individuals (excluding AB and C13E) (top) and GnRH control

(bottom)
