## supplementary table for "Generation of the First Transgenic Line of the Iconic Coral Reef Fish *Amphiprion ocellaris*"

| **(A)** | **Parenting vs Artificial Incubation** | | |
| --- | --- | --- | --- |
| *Predictors* | *Odds Ratios* | *CI* | *p* |
| (Intercept) | 18.83 | 12.71 – 27.90 | **<0.001** |
| Condition [Injected] | 0.22 | 0.18 – 0.28 | **<0.001** |
| Condition [Parenting] | 12.21 | 4.38 – 34.06 | **<0.001** |
| Dpf [2] | 0.07 | 0.07 – 0.08 | **<0.001** |
| Dpf [3] | 0.04 | 0.04 – 0.04 | **<0.001** |
| Dpf [4] | 0.03 | 0.02 – 0.03 | **<0.001** |
| Dpf [5] | 0.02 | 0.02 – 0.02 | **<0.001** |
| Dpf [6] | 0.01 | 0.01 – 0.02 | **<0.001** |
| Dpf [7] | 0.03 | 0.02 – 0.04 | **<0.001** |
| **Random Effects** | | | |
| σ^2^ | 3.29 | | |
| τ_00_ _Tile:(Batch:Animal)_ | 0.69 | | |
| τ_00_ _Batch:Animal_ | 0.39 | | |
| τ_00_ _Animal_ | 0.31 | | |
| ICC | 0.30 | | |
| N _Tile_ | 160 | | |
| N _Batch_ | 47 | | |
| N _Animal_ | 21 | | |
| Observations | 957 | | |
| Marginal R^2^ / Conditional R^2^ | 0.377 / 0.563 | | |

**(B)**

| Contrast | Odds.ratio | SE | Df | Asymp.LCL | Asymp.UCL | Z.ratio | P.Value |
| --- | --- | --- | --- | --- | --- | --- | --- |
| Ctrl/Injected | 4.4866 | 0.5212 | Inf | 3.57306 | 5.6337 | 12.923 | <.0001 |
| Ctrl/Parenting | 0.0819 | 0.0429 | Inf | 0.02936 | 0.2285 | -4.781 | <.0001 |
| Injected/Parenting | 0.0183 | 0.0095 | Inf | 0.00659 | 0.0506 | -7.697 | <.0001 |

**Supplementary Table 1. A)** Glmer estimates used to compare eggs reared under artificial incubation to eggs reared by parents. **B)** Post-hoc pairwise comparisons for survival across the three measured conditions calculated using “emmeans”.

|  | **Injected Eggs + Sterilization** | | |
| --- | --- | --- | --- |
| *Predictors* | *Odds Ratios* | *CI* | *p* |
| (Intercept) | 11.64 | 2.91 – 46.52 | **0.001** |
| Treatment [Fresh] | 1.38 | 0.44 – 4.30 | 0.577 |
| Treatment [H2O2] | 2.32 | 1.02 – 5.28 | **0.044** |
| Treatment [Iodine] | 1.18 | 0.62 – 2.26 | 0.611 |
| Treatment [KCL] | 3.49 | 1.62 – 7.50 | **0.001** |
| Treatment [MB] | 1.55 | 0.77 – 3.12 | 0.223 |
| Sterilization [UV] | 1.68 | 0.62 – 4.56 | 0.305 |
| Needle | 0.86 | 0.79 – 0.94 | **0.001** |
| Dpf [2] | 0.06 | 0.05 – 0.07 | **<0.001** |
| Dpf [3] | 0.03 | 0.03 – 0.03 | **<0.001** |
| Dpf [4] | 0.02 | 0.01 – 0.02 | **<0.001** |
| Dpf [5] | 0.01 | 0.01 – 0.01 | **<0.001** |
| Dpf [6] | 0.01 | 0.01 – 0.01 | **<0.001** |
| **Random Effects** | | | |
| σ^2^ | 3.29 | | |
| τ_00_ _Tile:(Batch:Animal)_ | 0.43 | | |
| τ_00_ _Batch:Animal_ | 0.27 | | |
| τ_00_ _Animal_ | 0.24 | | |
| ICC | 0.22 | | |
| N _Tile_ | 95 | | |
| N _Batch_ | 44 | | |
| N _Animal_ | 18 | | |
| Observations | 548 | | |
| Marginal R^2^ / Conditional R^2^ | 0.396 / 0.532 | | |

**Supplementary Table 2.** Glmer estimates used to establish effects of sterilization methods, dips, and needle diameter on survival for injected eggs.

|  | **Injected Eggs + Reagent Concentrations** | | |
| --- | --- | --- | --- |
| *Predictors* | *Odds Ratios* | *CI* | *p* |
| (Intercept) | 111.31 | 22.55 – 549.38 | **<0.001** |
| Needle | 0.87 | 0.72 – 1.05 | 0.157 |
| Ef1α Conc. | 0.98 | 0.95 – 1.00 | 0.102 |
| *Tol2* Conc. | 0.94 | 0.90 – 0.99 | **0.015** |
| Ef1α X *Tol2* | 1.00 | 1.00 – 1.00 | 0.055 |
| Dpf [2] | 0.05 | 0.05 – 0.06 | **<0.001** |
| Dpf [3] | 0.03 | 0.02 – 0.03 | **<0.001** |
| Dpf [4] | 0.01 | 0.01 – 0.01 | **<0.001** |
| Dpf [5] | 0.01 | 0.01 – 0.01 | **<0.001** |
| Dpf [6] | 0.01 | 0.01 – 0.01 | **<0.001** |
| **Random Effects** | | | |
| σ^2^ | 3.29 | | |
| τ_00_ _Tile:(Batch:Animal)_ | 0.53 | | |
| τ_00_ _Batch:Animal_ | 0.36 | | |
| τ_00_ _Animal_ | 0.00 | | |
| ICC | 0.21 | | |
| N _Tile_ | 44 | | |
| N _Batch_ | 20 | | |
| N _Animal_ | 13 | | |
| Observations | 260 | | |
| Marginal R^2^ / Conditional R^2^ | 0.436 0/556 | | |

**Supplementary Table 3.** Glmer estimates used to establish effects of transgene reagents on survival for injected eggs.

|  | **Injected Eggs + Time to Inject** | | |
| --- | --- | --- | --- |
| *Predictors* | *Odds Ratios* | *CI* | *p* |
| (Intercept) | 96.02 | 5.96 – 1547.85 | **0.001** |
| Needle | 0.69 | 0.54 – 0.87 | **0.002** |
| Time to Inject | 1.02 | 1.01 – 1.03 | **0.008** |
| Ef1α Conc. | 1.02 | 1.00 – 1.05 | 0.060 |
| *Tol2* Conc. | 0.93 | 0.82 – 1.05 | 0.249 |
| Dpf [2] | 0.08 | 0.06 – 0.10 | **<0.001** |
| Dpf [3] | 0.04 | 0.03 – 0.05 | **<0.001** |
| Dpf [4] | 0.02 | 0.01 – 0.02 | **<0.001** |
| Dpf [5] | 0.01 | 0.01 – 0.01 | **<0.001** |
| Dpf [6] | 0.01 | 0.01 – 0.01 | **<0.001** |
| **Random Effects** | | | |
| σ^2^ | 3.29 | | |
| τ_00_ _Tile:(Batch:Animal)_ | 0.21 | | |
| τ_00_ _Batch:Animal_ | 0.00 | | |
| τ_00_ _Animal_ | 0.00 | | |
| ICC | 0.06 | | |
| N _Tile_ | 15 | | |
| N _Batch_ | 5 | | |
| N _Animal_ | 5 | | |
| Observations | 89 | | |
| Marginal R^2^ / Conditional R^2^ | 0.475 / 0.507 | | |

**Supplementary Table 4.** Glmer estimates used to establish effect of time to inject on survival for injected eggs.

|  | **Control Eggs + Time to Take** | | |
| --- | --- | --- | --- |
| *Predictors* | *Odds Ratios* | *CI* | *p* |
| (Intercept) | 7.33 | 0.87 – 62.06 | 0.068 |
| Treatment [Fresh] | 0.85 | 0.31 – 2.37 | 0.760 |
| Treatment [H2O2] | 1.86 | 0.53 – 6.55 | 0.331 |
| Treatment [Iodine] | 1.12 | 0.24 – 5.25 | 0.884 |
| Treatment [KCL] | 1.15 | 0.16 – 8.31 | 0.892 |
| Treatment [MB] | 0.44 | 0.09 – 2.14 | 0.312 |
| Time to Take | 1.08 | 0.98 – 1.19 | 0.111 |
| Dpf [2] | 0.05 | 0.04 – 0.06 | **<0.001** |
| Dpf [3] | 0.02 | 0.02 – 0.03 | **<0.001** |
| Dpf [4] | 0.01 | 0.01 – 0.02 | **<0.001** |
| Dpf [5] | 0.01 | 0.01 – 0.01 | **<0.001** |
| Dpf [6] | 0.01 | 0.01 – 0.01 | **<0.001** |
| **Random Effects** | | | |
| σ^2^ | 3.29 | | |
| τ_00_ _Tile:(Batch:Animal)_ | 0.40 | | |
| τ_00_ _Batch:Animal_ | 0.88 | | |
| τ_00_ _Animal_ | 0.17 | | |
| ICC | 0.31 | | |
| N _Tile_ | 27 | | |
| N _Batch_ | 17 | | |
| N _Animal_ | 11 | | |
| Observations | 160 | | |
| Marginal R^2^ / Conditional R^2^ | 0.387 / 0.575 | | |

**Supplementary Table 5.** Glmer estimates used to establish effect of time to take on survival for control eggs.

|  | **Control Eggs + Sterilization** | | |
| --- | --- | --- | --- |
| *Predictors* | *Odds Ratios* | *CI* | *p* |
| (Intercept) | 3.85 | 0.46 – 32.12 | 0.213 |
| Treatment [Fresh] | 0.83 | 0.32 – 2.16 | 0.703 |
| Treatment [H2O2] | 1.59 | 0.61 – 4.11 | 0.343 |
| Treatment [Iodine] | 0.82 | 0.33 – 2.00 | 0.661 |
| Treatment [KCL] | 0.83 | 0.29 – 2.43 | 0.740 |
| Treatment [MB] | 0.59 | 0.18 – 1.92 | 0.378 |
| Sterilization [UV] | 2.99 | 0.37 – 24.39 | 0.306 |
| Dpf [2] | 0.12 | 0.10 – 0.13 | **<0.001** |
| Dpf [3] | 0.07 | 0.06 – 0.08 | **<0.001** |
| Dpf [4] | 0.05 | 0.04 – 0.05 | **<0.001** |
| Dpf [5] | 0.04 | 0.03 – 0.04 | **<0.001** |
| Dpf [6] | 0.03 | 0.03 – 0.04 | **<0.001** |
| Dpf [7] | 0.05 | 0.03 – 0.07 | **<0.001** |
| **Random Effects** | | | |
| σ^2^ | 3.29 | | |
| τ_00_ _Tile:(Batch:Animal)_ | 0.79 | | |
| τ_00_ _Batch:Animal_ | 0.23 | | |
| τ_00_ _Animal_ | 0.49 | | |
| ICC | 0.31 | | |
| N _Tile_ | 65 | | |
| N _Batch_ | 41 | | |
| N _Animal_ | 18 | | |
| Observations | 383 | | |
| Marginal R^2^ / Conditional R^2^ | 0.230 / 0.472 | | |

**Supplementary Table 6.** Glmer estimates used to establish effects of sterilization methods and dips on survival for control eggs.

|  | **Rate of Transgenesis** | | |
| --- | --- | --- | --- |
| *Predictors* | *Odds Ratios* | *CI* | *p* |
| (Intercept) | 0.14 | 0.05 – 0.43 | **0.001** |
| Ef1α Conc. | 1.03 | 1.00 – 1.06 | 0.081 |
| *Tol2* Conc. | 1.08 | 1.02 – 1.14 | **0.010** |
| Ef1α X *Tol2* | 1.00 | 1.00 – 1.00 | 0.207 |
| **Random Effects** | | | |
| σ^2^ | 3.29 | | |
| τ_00_ _Animal_ | 0.52 | | |
| ICC | 0.14 | | |
| N _Animal_ | 15 | | |
| Observations | 54 | | |
| Marginal R^2^ / Conditional R^2^ | 0.116 / 0.235 | | |

**Supplementary Table 7.** Glmer estimates for effects of reagents on rate of transgenesis.
